# Htz1 and Set1 Regulate Ergosterol Levels in Response to Environmental Stress

**DOI:** 10.1101/199174

**Authors:** Kubra Aslan, Bilge Özaydin

## Abstract

Ergosterol is an essential isoprenoid for cellular integrity and proper membrane fluidity of fungi. Proper level of ergosterol is crucial for resistance to various stressful conditions, such as hypoxia, hypothermia, and hyperosmolarity. The isoprenoid building blocks of ergosterol are synthesized via the mevalonate pathway, which relies on the availability of many central metabolites, such as acetyl-coA and S-adenosyl methionine (SAM). The metabolic currencies are also the substrates for epigenetic modifications such as histone acetylation and methylation. To have a better understanding of how isoprenoid synthesis and these epigenetic mechanisms affect each other, we re-analyzed the results of our screen on *Saccharomyces cerevisiae* gene deletion collection for isoprenoid production and found a group of chromatin regulators with significant effects on isoprenoid production. More specifically, the canonical histone Htz1 (H2A.z), the SWR1 complex that loads Htz1 onto chromatin, and the histone demethylase Jhd2 inhibited, whereas the Htz1 unloading INO80 complex and histone methylase Set1 enhanced isoprenoid production. Further analysis of genome-wide expression data revealed that Htz1 and Set1 differentially regulate stress-response genes which presumably affect isoprenoid synthesis. Conversely, changes in isoprenoid production alters the transcription of the same set of genes. Further analysis of ergosterol levels in these gene deletions showed that *htz1, set1* double deletion leads to accumulation of ergosterol beyond homeostatic levels and renders cells vulnerable to environmental stress. Our re-analysis of multiple published data and follow-up experiments revealed an epigenetic crosstalk mechanism between ergosterol levels and stress response genes that is essential for maintaining optimum concentration of ergosterol under various conditions.

## INTRODUCTION

Isoprenoids are terpene derived organic compounds found in almost all organisms (Holstein and Raymond, 2004). In addition to their biological functions, many isoprenoids have extensive use in agricultural, pharmaceutical, and food industries (Ro *et al.*, 2006, Ajikumar *et al.*, 2010). These commercially valuable isoprenoids are often secondary metabolites that are produced in limited and stochastic quantities in their natural resources. Metabolic engineering of microbial organisms provides a sustainable, cost-effective, and fast means to producing high yields of pure form of desired isoprenoid compounds (Vickers *et al.*, 2017). *Saccharomyces cerevisiae*, the budding yeast, is often the organism of choice for such processes with its established molecular biology techniques and long history of industrial applications (Hong and Nielsen, 2012; Paramasiyan and Mutturi, 2017).

In yeast, isoprenoids are synthesized via the mevalonate pathway, which condenses three acetyl-coA molecules to 5-carbon isoprene compounds that serve as the building clocks for production of various isoprenoids including ergosterol, the fungal sterol with roles in membrane fluidity and integrity. Synthesis of a single molecule of ergosterol requires 18 molecules of acetyl-coA, 17 molecules of NADPH, 12 molecules of ATP, and single molecule of S-adenosyl-methionine (SAM), making it metabolically very costly (Parks and Casey, 1995). Each one of these intermediary metabolites are not only utilized in biosynthetic pathways but they are also the substrates for various histone modifications that play role in the transcriptional regulation of the genome (Janke *et al.*, 2015; Etchegaray and Mostoslavsky, 2016). Intracellular concentrations of these metabolites are crucial determinants for both the flux of the metabolic pathways and the level of histone modifications. For example, acetyl-coA levels are directly influenced by nutrient availability and multiple studies showed cellular acetyl-coA levels correlate with histone acetylation levels (Zhang *et al*., 2013; Galdieri *et al*., 2013; Shi and Tu, 2015). S-adenosyl methionine (SAM) is another metabolite that is crucial for synthesis of various isoprenoids and also important for gene regulation via histone methylation. When ergosterol synthesis is down-regulated, Sam levels increased threefold, suggesting that ergosterol synthesis is a significant competitor for intracellular SAM pools (Shobayashi *et al*., 2006). SAM is also the substrate for histone methylation at lysine residues of histones H3K4, H3K36, and HK79 (Sadhu *et al*. 2013). These methylation sites act as a docking site for chromodomain-containing transcription regulators and can result in either repression or activation of the genes they are found at. Therefore these metabolites create a link between ergosterol metabolism and transcriptional regulation of genome via histone modifications.

Our previous work (Özaydin *et al.*, 2013) used carotenogenic plasmids to phenotypically screen the haploid yeast deletion collection for changes in isoprenoid production. In this paper, we re-analyzed the data from this screen and other published work for various chromatin regulatory factors and revealed that ergosterol levels are regulated by combinatorial actions of the canonical histone Htz1 and the histone methyl transferase Set1. Set1 dependent repression of stress response genes resulted in decreased ergosterol production, whereas Htz1 and Jhd2 dependent activation of the same subset of genes increased ergosterol production. When both of these regulatory mechanisms were abolished, cells accumulated ergosterol beyond homeostatic levels and lost the ability to resist various stressful conditions. This work, for the first time, shows a cross-talk mechanism between ergosterol levels and environmental stress conditions that is mediated through multiple chromatin regulatory complexes.

## MATERIALS AND METHODS

### Yeast strains, media and transformation

Yeast strains used in this study were isogenic to BY4741 and listed in Table 1.

**Table 1.** Strains used in this study.

| Strain name | Accession number | Genotype | Source |
| --- | --- | --- | --- |
| wt (BY4741) | Y00000 | MATa; his3Δ1; leu2Δ0; met15Δ0; ura3Δ0 | Euroscarf |
| *htz1* | Y01703 | BY4741; YOL012c::kanMX4 | Euroscarf |
| *ies2* | Y01997 | BY4741; YNL215w::kanMX4 | Euroscarf |
| *jhd2* | Y06922 | BY4741; YJR119c::kanMX4 | Euroscarf |
| *set1* | SDBY1210 | BY4741; YHR119w::hygMX | Scott Briggs |
| *htz1, set1* | YBO087 | SDBY1210; YOL012c::kanMX4 | This study |

Rich medium (YPD) is described previously (Sherman *et al.*, 1999). KanMX4 resistance marker was selected on YPD containing 200 mg/liter of G418 (Geneticin). Modified lithium acetate transformation was used as described previously (Becker and Lundblad, 2001).

Gene deletions and genomic integrations were done using one-step integration of PCR-amplified knockout cassettes (Goldstein and McCusker, 1999; Longtine et al., 1998) and confirmed by PCR and phenotypic validation.

Oligonucleotide sets used for *htz1* deletion and colony PCR verification are in Table 2

**Table 2.**
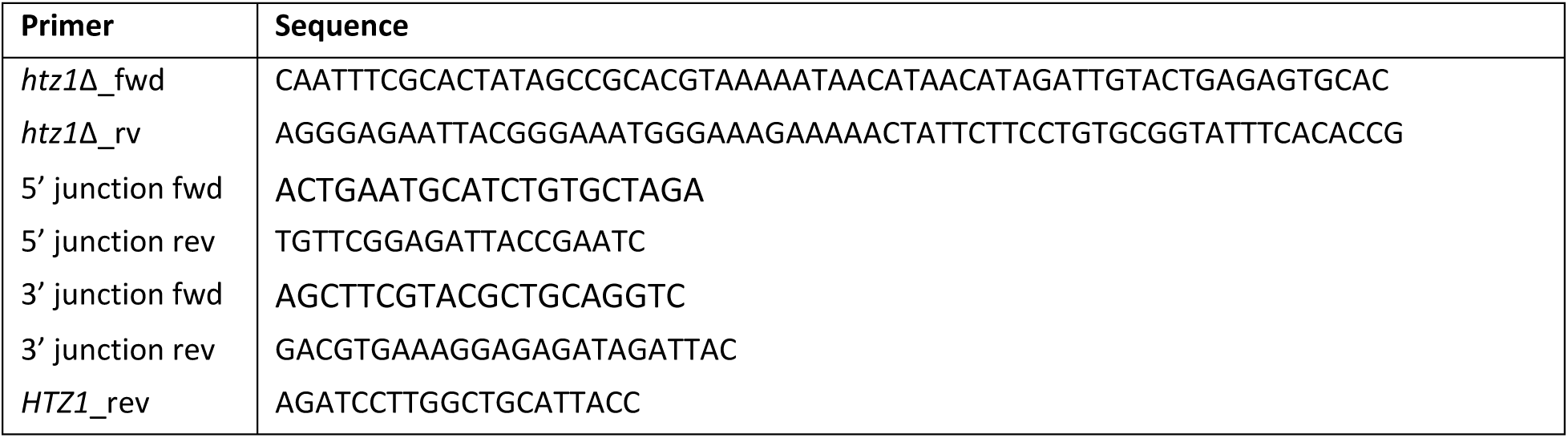
Oligonucleotides used in this study.

### Extraction and analysis of ergosterol

Total sterol was isolated from whole yeast cells by saponification, followed by extraction of non-saponifiable lipids with hexane (Breivik and Owade, 1957). 20 ml of yeast culture was harvested and washed twice with water. Cell pellets were lyophilized and dry cell mass was determined. Dried cells were resuspended in 4 ml of 25% alcoholic KOH solution (25g of KOH in 70% ethanol) and incubated at 80 ^o^C for 1 hour by vortexing every 10 minutes. Sterols were extracted by adding equal volume of hexane and vortexing for 1 minute. After spinning at 2800 rpm, the upper layer was retained and absorption was measured in the 240nm-330nm spectrum. The presence of ergosterol and intermediate 24(28)dehydroergosterol showed the characteristic four-peaked curve in this spectrum.

Total ergosterol content was calculated as a percentage of dry cell weight (dcw) using the following equations where F is the factor for dilution in hexane and 290 and 518 are the E values (in percentages per centimeter) determined for crystalline ergosterol and 24(28)DHE, respectively.

% ergosterol + % 24(28)DHE = [(A_281.5_/290) × F] /dcw

% 24(28)DHE = [(A_230_/518) × F] /dcw

% ergosterol = [% ergosterol + % 24(28)DHE] - % 24(28)DHE

### Iodine staining

10-fold serial dilutions of overnight yeast cultures were spotted on rich media and were grown for 3 days. Iodine crystals were poured to cover the lid of the petri dish and yeast colonies were exposed to iodine vapor until they developed color, for about 2-3 minutes (Chester, 1968).

### Gene ontology (GO) term classification

A list of chromatin regulatory complexes was formed based on GO term classification. We analyzed the microarray data on gene deletions for differentially regulated pathways and processes by searching for over-represented gene ontology (GO) terms. Significantly enriched gene ontology terms (p < 0.01) were clustered by GO slim macromolecular complex terms using GO term mapper (http://go.princeton.edu/cgi-bin/GOTermMapper). If more than one GO term in the same “family” were identified, only the most significant term was listed.

## RESULTS

### Chromatin remodeling complexes SWR1C and INO80C differentially affect isoprenoid production

Previous screen of the yeast haploid deletion collection with carotenogenic genes (Özaydin *et al.*, 2013) characterized more than 300 genes with significant effects on the isoprenoid production. In this study, we focused on the chromatin regulator genes to have a better understanding of how epigenetic factors regulate isoprenoid synthesis. A list of chromatin regulatory complexes was formed using GO slim macromolecular complex terms (Supplementary table 1). Carotenoid production of each deletion strain is scored at a scale of -5 to 5, where -5 is white, 0 is the color of the wild-type parent strain, and 5 is the darkest orange color. The color scores of the deletions of the members of each chromatin regulatory complex are plotted using the screen results (Figure 1A). Complexes with more than half of the members that were not scored are excluded from this plot. The median color score of most complexes were close to zero, indicating no significant effect on carotenoid production. However, members of two complexes, SWR1 and INO80, had the most concordant and significant effect on the carotenoid production with median color scores of 3 and -2.5 respectively. Both SWR1C and INO80C are ATP-dependent chromatin regulators that load and unload the histone variant H2A.z (Htz1) respectively (Krogan *et al.*, 2003; Papamichos-Chronakis *et al.*, 2011). This analysis suggested that SWR1 complex dependent binding of Htz1 to the chromatin decreased, whereas its removal by INO80 complex enhanced carotenoid production. In accordance with this conclusion, deletion of *HTZ1* increased carotenoid production (Table 3).

**Table 3.** Chromatin regulators with the highest effect on carotenoid production (color score ≥4 and ≤-4)

| Chromatin regulators | Function | Deletion score |
| --- | --- | --- |
| ARP6 | Actin-related protein; a component of the SWR1 complex. | 5 |
| SWR1 | Swi2/Snf2-related ATPase; structural component of the SWR1 complex. | 4 |
| YAF9 | Subunit of NuA4 histone H4 acetyltransferase and SWR1 complexes.. | 4 |
| HTZ1 | Histone variant H2AZ; exchanged for histone H2A in nucleosomes by the SWR1 complex | 4 |
| JHD2 | JmjC domain family histone demethylase; promotes global demethylation of H3K4. | 5 |
| INO80 | ATPase and nucleosome spacing factor; unloads Htz1 from chromatin. | -4 |
| IES2 | Associates with INO80 chromatin remodeling complex. | -4 |
| SUS1 | Member of SAGA histone acetylase and TREX-2 complexes, mRNA export coupled transcription activation and elongation. | -5 |
| LDB7 | Component of the RSC chromatin remodeling complex. | -5 |
| PAF1 | Transcription elongation, required for expression and repression of a subset of genes | -4 |
| SET1 | Histone methyltransferase for H3K4, subunit of the COMPASS (Set1C) complex. | Not available |

**Figure 1.**
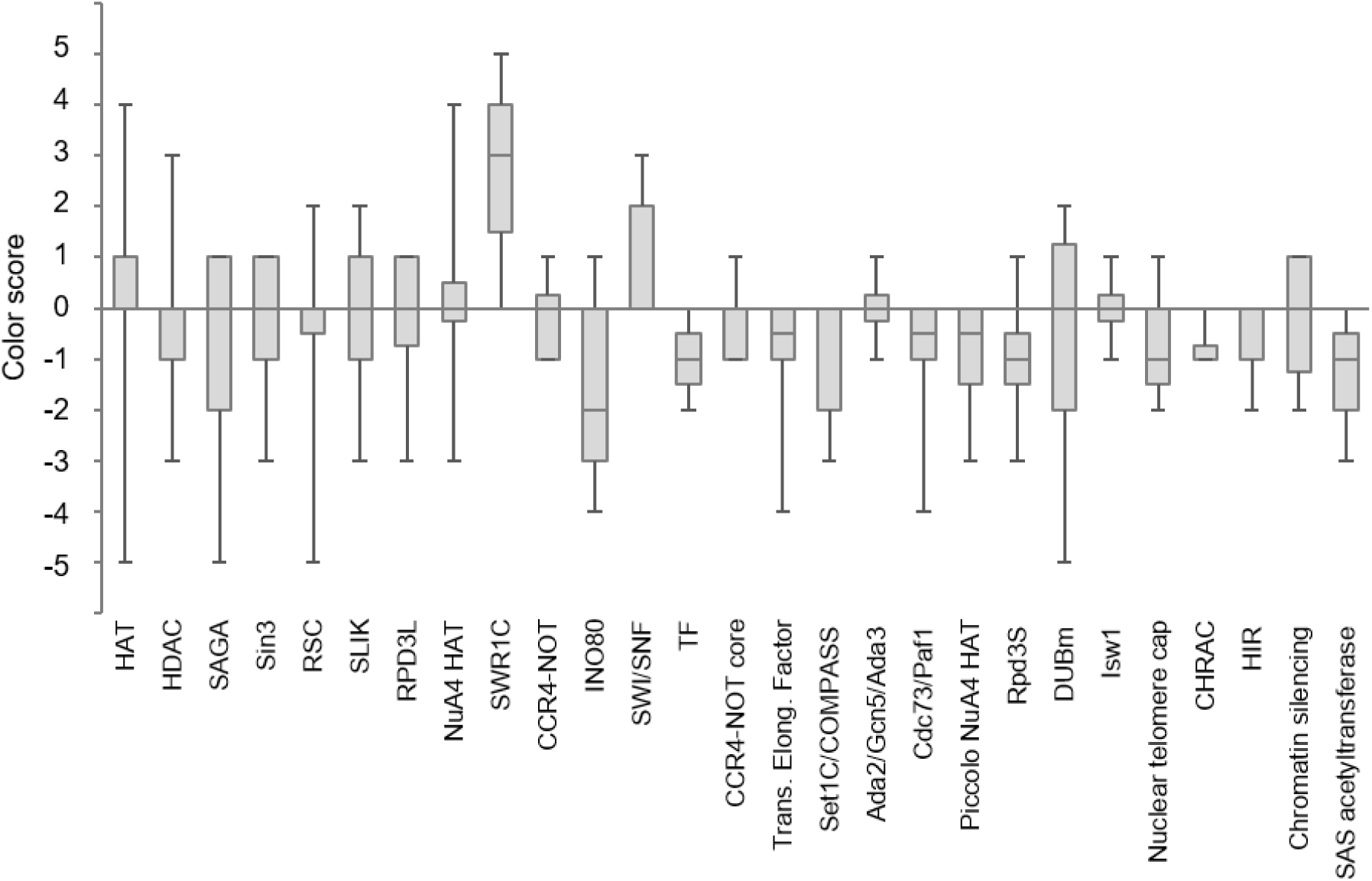
Color score distribution of the deletion of chromatin regulator complexes. Names of the chromatin regulatory complexes are indicated on the x-axis. Whiskers indicate the maximum and minimum color scores of the respective complex member, the line inside the box indicates the median of the scores.

### Set1 dependent methylation of Histone 3 lysine 4 (H3K4) is a significant epigenetic mark for isoprenoid biosynthesis

To elucidate the epigenetic mechanisms affecting isoprenoid production, we focused on the individual chromatin regulators with the highest effect on isoprenoid production (Table 3). In addition to the members of SWR1 and INO80 complexes, only a handful of genes (*JHD2*, *SUS1*, *LDB7*, *PAF1*) altered isoprenoid production significantly. Jhd2 is the only chromatin regulator besides SWR1 complex members that increased carotenoid production when deleted. Jhd2 is a jumonji domain histone demethylase that removes the tri-methyl group on the fourth lysine residue of histone H3 (H3K4me^3^) that is introduced by the methyl transferase Set1. Set1 deletion strain was missing from the yeast haploid deletion collection, therefore its effect on carotenoid production was not determined. However, Set1 is a member of COMPASS complex (Krogan et *al.*, 2002) and deletion of different subunits of COMPASS decreased carotenoid production (Figure 1). Set1 was also recently shown to enhance ergosterol production through transcriptional regulation of multiple genes in the mevalonate pathway (South *et al.,* 2013). Together, these data suggest that H3K4me^3^ is an important epigenetic mark that effects ergosterol production as well as production of other isoprenoids.

### Set1 and Htz1 differentially regulate genes with roles in cell wall biosynthesis and stress response

Genome-wide expression patterns of the deletions of chromatin regulators listed in Table 3 were compared to reveal the genomic perturbations they caused and thereby find clues to how they may affect the isoprenoid synthesis. Recent work by Kemmeren *et al.* (2014) examined genome-wide mRNA expression changes in individual deletions of one-quarter of yeast genes. Except for *ino80*, all the genes listed in Table 3 were included in this study (Figure 2A). Clustering analysis revealed two distinct groups of chromatin regulators with opposite effects on the genome, except for *ldb7* deletion, which had a more distinctive expression profile. Deletion of SWR1 complex members and *htz1* showed similar expression profiles with mostly down-regulation of gene expression, except for a small cluster of genes with roles in methionine biosynthesis and arginine degradation. Deletions of *ies2*, *set1*, and RNA polymerase-II associated factors *sus1* and *paf1* formed another group with similar expression profiles. Paf1 is a transcription elongation factor that is recruited to chromatin through Set1 dependent histone methylation (Mulder *et al.*, 2007), which explains the concordance in expression profiles.

**Figure 2.**
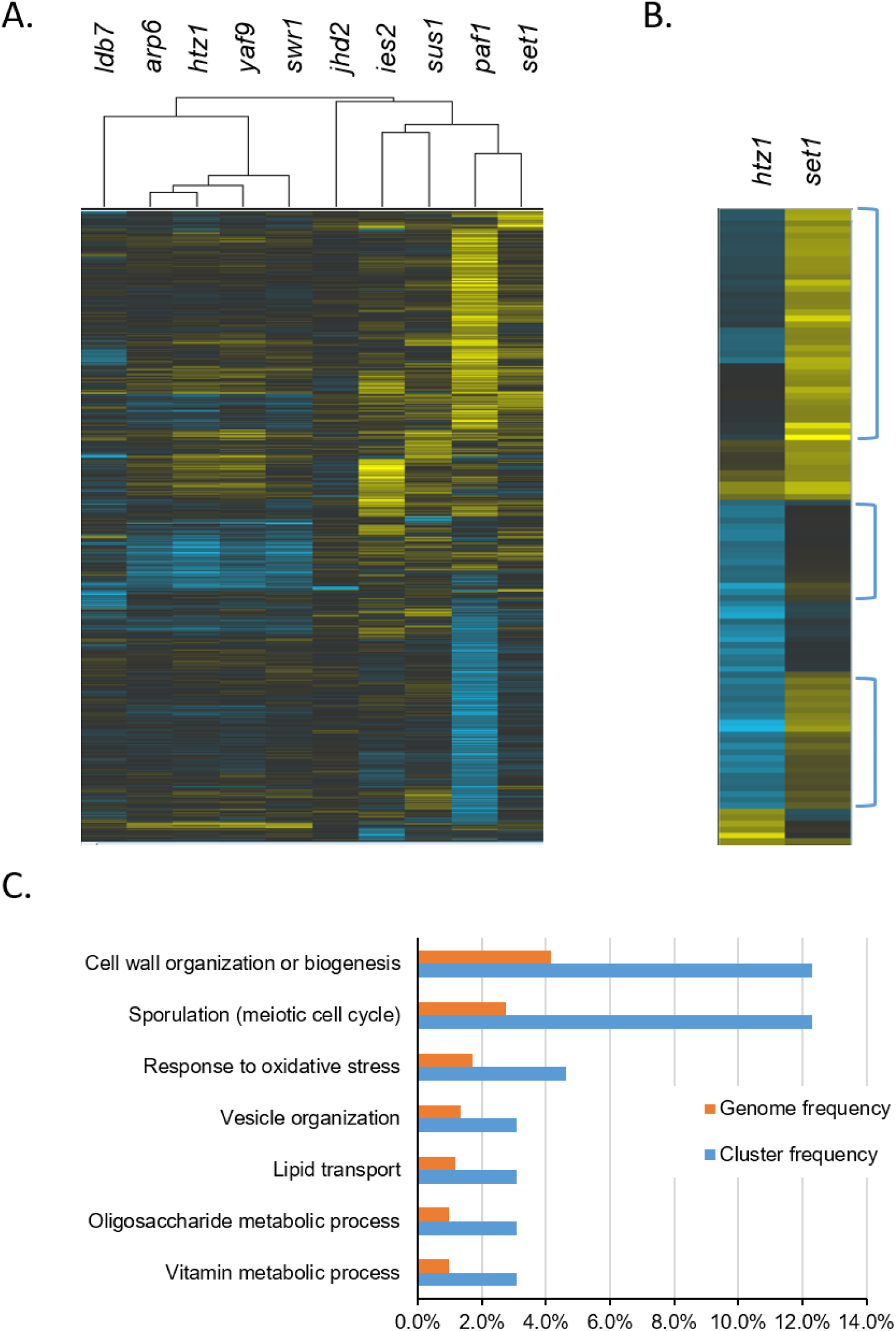
A. Clustering of transcriptional profiles of single gene deletions (**Kemmeren *et al.*, 2014**) with highest effect on carotenoid production (<-3 and >3) **B.** Comparing the transcriptional profile of the deletions of major CR players, htz1 and set. **C.** Gene ontology (GO) term mapping of those genes differentially regulated between htz1 and set1 strains.

Sus1 (Pascual-García and Rodríguez-Navarro, 2009) and Paf1 are downstream regulators of transcription, whereas Htz1 loading and Set1 methylation dictate which genes to be transcribed or not. Therefore, we compared genome-wide expression profiles of *set1* and *htz1* as the upstream players of the gene regulation to have a clearer picture of the transcriptional regulation by these two groups (Figure 2B). The most differentially expressed genes between *set1* and *htz1* as indicated with brackets in Figure 2B (See Supplementary Table 2 for the list of the genes) were analyzed using gene ontology term mapper. Sporulation and cell wall biogenesis related genes were four-times and oxidative stress response genes and lipid and oligosaccharide synthesis related genes were three-times more enriched in this cluster of genes than the genome. Ergosterol and other isoprene derived molecules such as heme and dolichol play central roles in all of these biological processes (Deng *et al.*, 2008; Liu *et al.*, 2017, Grabinska *et al.*, 2008). Therefore, *set1* and *htz1* dependent gene expression changes had implications for the isoprenoid metabolism.

Genes that were induced or repressed more than 1.5-fold in *htz1* and *set1* are mapped onto the chromosome (Figure 3). Of the 110 genes that were up-regulated in *set1*, 26 of them were down-regulated in *htz1* (Table 2A). When 10kb neighborhood of these 110 genes were scanned, *htz1* down-regulated genes were found in 50 of them. On the other hand, almost all of the genes that were down-regulated in *set1* was unchanged in *htz1*. This analysis suggested that genes activated by Set1 were not affected by Htz1, but about half of the Set1 inhibited genes were activated in an Htz1-dependent manner. Therefore, genome-wide expression data for *htz1* and *set1* strains indicated a strong correlation for Htz1 dependent activation of Set1 repressed genes (p<10^-6^).

**Figure 3.**
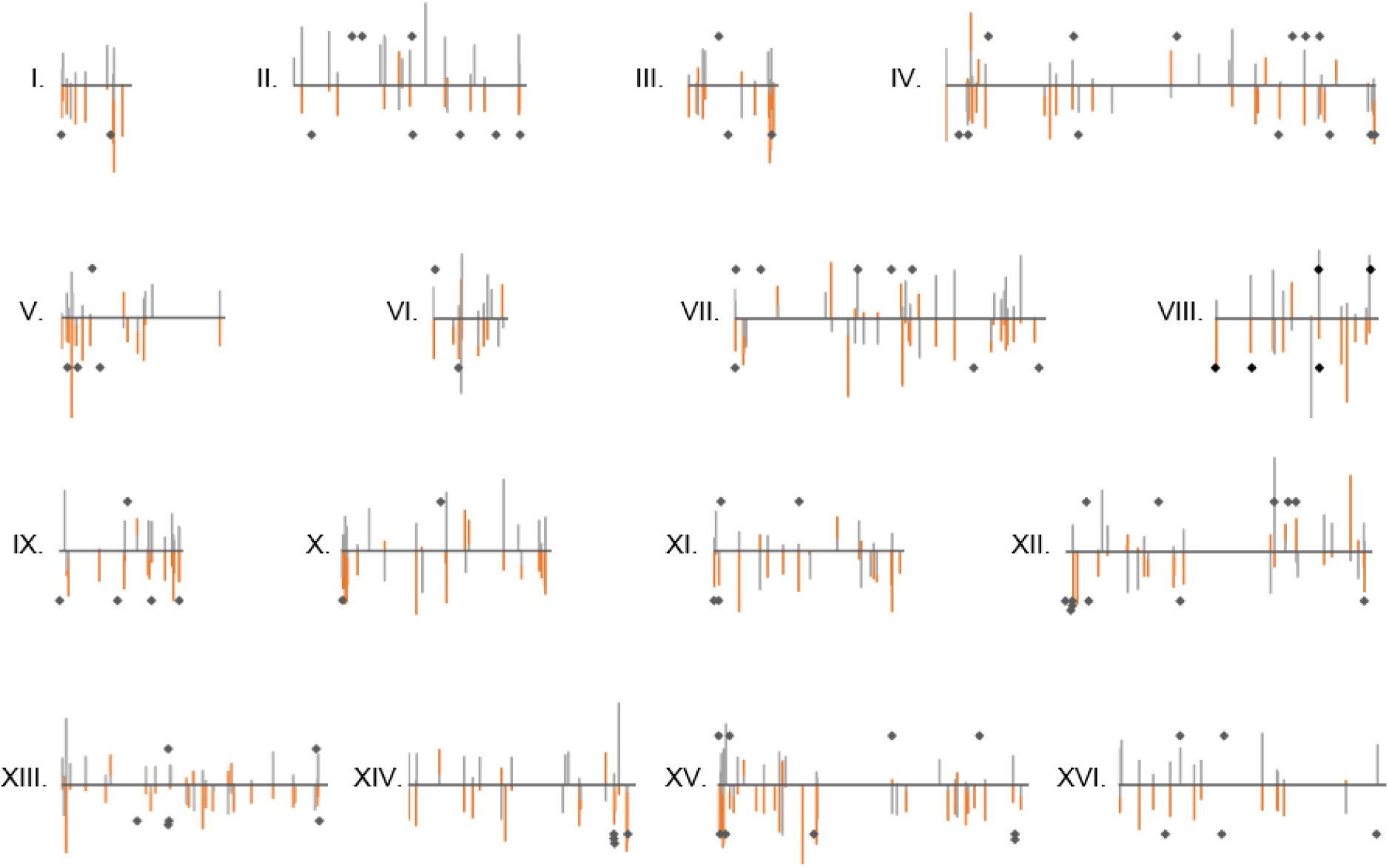
Gene perturbations in *set1* (gray lines) and *htz1* (red lines) strains mapped on chromosomes (Kemmeren *et al.*, 2014). Black diamonds are genes upregulated or downregulated upon carotenoid over-production (Verwaal *et al.*, 2010).

### Over-production of carotenoids results in transcriptional changes in genes that are regulated by Htz1 and Set1

Verwaal *et al.* (2010) showed that high-level production of carotenoids represses 57 genes and induces 36 genes, among which many stress response proteins are found. Our analysis showed that, about 80% of these genes are either directly affected by *set1* and/or *htz1* deletions or are in close proximity (<10kb) of genes affected by *htz1* and *set1* (Figure 3, black diamonds on the graph). Of the 57 repressed genes found in carotenoid producing yeast, 42 are also found in Htz1 activated regions and 40 are in Set1 repressed domains (Table 4). More than half of the 36 induced genes are also up-regulated in *set1* strain and down-regulated in *htz1* strain. This analysis suggested that changes in isoprenoid production induced genome-wide transcriptional changes through Set1 and Htz1 dependent mechanisms.

**Table 4.** Number of genes that are regulated in *htz1* and *set1* strains

|  | *set1*<br>high | *set1*<br>low | *set1*<br>unchanged |  |
| --- | --- | --- | --- | --- |
| *htz1* high | 0 | 0 | 0 | Genes down-regulated in carotenoid<br>producing yeast |
| *htz1* low | 37 | 0 | 6 |  |
| *htz1*<br>unchanged | 4 | 0 | 10 |  |
| *htz1* high | 0 | 2 | 0 | Genes up-regulated in carotenoid<br>producing yeast |
| *htz1* low | 16 | 0 | 4 |  |
| *htz1*<br>unchanged | 3 | 1 | 9 |  |

### Htz1 and Set1 Regulates Ergosterol Levels

Synthesis of both carotenoids and ergosterol compete for the same isoprene building blocks, FPP and GPP. Therefore, the transcriptional changes observed in carotenoid over-producing strain and *htz1* strain (Figure 3) may down-regulate ergosterol synthesis which in turn will result in increased carotenoid production. The fact that *htz1* shares the most phenotypic properties with the deletion of sterol methyltransferase *erg6*, a determining step for ergosterol biosynthesis, also suggests that *htz1* results in decreased ergosterol levels. (Figure 4A).

**Figure 4.**
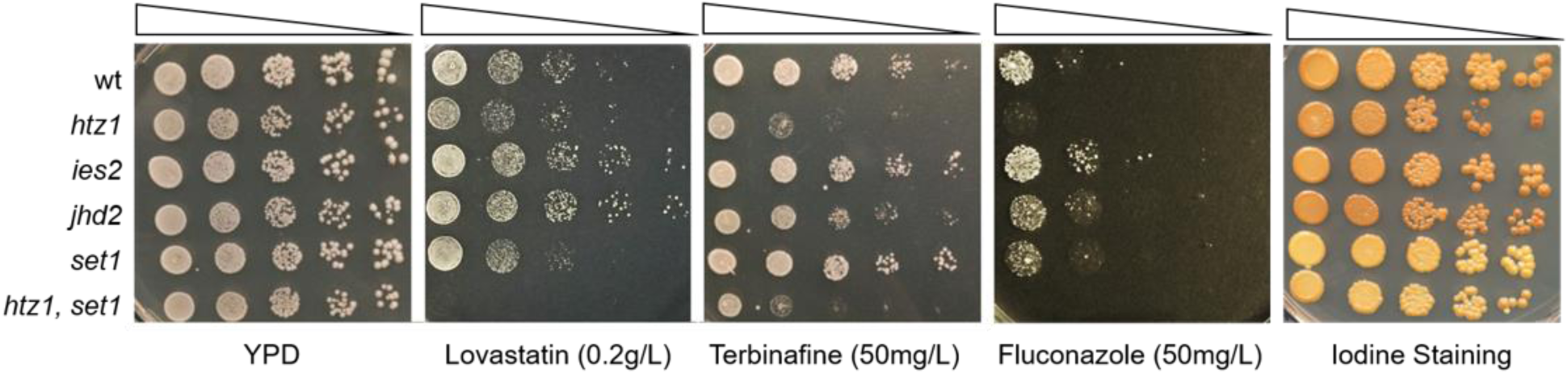
Drug sensitivity and glycogen storage levels in wild-type yeast strains and deletion strains *htz1*, *ies2*, *jhd2*, *set1*, and *htz1, set1*. Overnight cultures grown in YPD were spotted in 10-fold serial dilutions onto YPD plates, as well as YPD plates containing either lovastatin (0.2g/L), Terbinafine (50mh/L), or Fluconazole (50mg/L) drugs. Iodine staining is done by exposing YPD grown spots to iodine vapor.

To see whether the mevalonate or ergosterol pathways were affected in strains deleted for the characterized chromatin regulators, we tested their growth in the presence of three kinds of drugs that inhibit these pathways at different steps (Figure 4B). Lovastatin inhibits the HMGR enzyme, resulting in early inhibition of the mevalonate pathway and synthesis of all isoprene-derived molecules (Louis-Flamberg *et al.*, 1990). On the other hand, terbinafine and fluconazole only impedes ergosterol synthesis by inhibiting the first dedicated step of ergosterol synthesis and the demethylase enzyme that converts lanosterol to ergosterol respectively (Nowosielski *et al.*, 2011; Kelly *et al.*, 1997). Wild type strain and the deletion strains were spotted in 10-fold serial dilution on YPD plates containing lovastatin, terbinafine, or fluconazole. Deletions of *htz1* and most members of SWR1 complex were sensitive, whereas deletion of *ies2* and other members of INO80 complex were resistant to all the drugs tested (Figure 4B and Supplementary Figure 1). On the other hand, *set1* deletion was resistant to terbinafine but sensitive to lovastatin and fluconazole. Deletion of *jhd2* showed opposite effects to *set1*, as expected from its demethylase activity on Set1 methylated H3K4. The drug assay suggested that Htz1 upregulated the ergosterol synthesis, whereas Set1 inhibited it. Surprisingly, *set1, htz1* double deletion was the most sensitive strain to all the drugs tested, suggesting decreased production of both mevalonate and ergosterol.

One consequence of low ergosterol synthesis is the increase in the glycogen storage. We qualitatively measured glycogen levels by exposing cells to iodine vapor. As expected from the decrease in ergosterol synthesis, both *htz1* and *jhd2* showed increased glycogen staining. The set1 strain showed decreased glycogen storage in accordance with increased terbinafine resistance. However, the *set1, htz1* strain also showed very low glycogen levels despite drug assays indicating low level ergosterol synthesis in this double mutant. This suggested that *set1, htz1* strain could not upregulate ergosterol levels to counter balance the inhibitory drugs but stored ergosterol beyond homeostatic levels.

To measure ergosterol levels more directly, we extracted total sterols from yeast cells grown for 1-3 days and determined ergosterol levels using spectrophotometry. On the first day of growth, total ergosterol levels did not change significantly for *htz1* and *set1* but slightly decreased in *jhd2* and *ies1*. As the cells reached the late stationary phase (day 3), total ergosterol content increased almost two-fold in *set1, htz1* strain, while remaining same in other strains. This data was in accordance with the iodine staining, which showed decreased glycogen storage in *set1, htz1*, presumably as a result of increased ergosterol levels. Altogether, the data suggested that Htz1 and Set1 had opposing roles in maintaining the ergosterol homeostasis and deletion of both genes resulted in accumulation of ergosterol beyond homeostatic levels.

### Set1 and Htz1 Suppress Ergosterol Synthesis under Hyperosmotic Conditions

Ergosterol is down-regulated by a Hog1-dependent mechanism under hyperosmotic conditions (Montañés *et al.*, 2011). The stress-activated protein kinase, Hog1, activates stress response genes by either recruiting RSC complex to genes with methylated H3K4 or SWR1-C when H3K4 is un-methylated. This is in accordance with our hypothesis that Htz1 and Set1 maintain ergosterol homeostasis. To test whether Set1 and Htz1 is essential for down-regulating ergosterol under hyper-osmotic conditions, wild type and various yeast deletion strains were grown under normal (YPD) and hyper-osmotic (YPD + 1M NaCl) conditions. As expected, wild type yeast had almost two-fold less ergosterol levels under hyper-osmotic conditions (Figure 6A). Single gene deletions of *htz1*, *jhd2*, and *set1* only mildly decreased ergosterol levels. However, *set1, htz1* double knock-out strain not only did not fail to suppress ergosterol synthesis, but it accumulated two-fold more ergosterol than the wild-type strain. This data was in agreement of ergosterol increase in this strain under late stationary phase, which is supposedly a stressful condition for cells as well.

Inability to down-regulate ergosterol synthesis under hyper-osmotic conditions were shown to have negative effects on cell growth (Montañés *et al.*, 2011). To see if that was the case with the *set1, htz1* strain, deletion strains were spotted on YPD and YPD + 1M NaCl agar plates in 10-fold dilutions (Figure 6B). Wild type and single deletion strains grew equally well on YPD and YPD + 1M NaCl plates. On the other hand, the *htz1, set1* strain grew two orders of magnitude less under hyper-osmotic conditions, in agreement with the inability to suppress ergosterol synthesis upon salt stress.

## DISCUSSION

Most metabolic engineering efforts focus on maintaining high flux through the pathway of interest by increasing pathway precursors or upregulating the expression of the pathway genes. However, these efforts rarely move beyond proof of concept experiments and only result in limited amounts of target products. This outcome is not so surprising since overproduction of pathway proteins and molecules competes for cellular resources and consequently puts cells under stress. To further complicate the matter, the building blocks of most metabolic pathways are also used for various epigenetic modifications, such as histone methylation and acetylation, that can have significant genome wide transcriptional effects. Therefore, strain engineers must be mindful of cellular defense mechanisms, such as activation of stress response pathways, and epigenetic mechanisms that can act against the engineered pathway. This work sheds some light onto these mechanisms that get activated in response to changes in the ergosterol levels.

Besides its role in cell membrane integrity and fluidity, ergosterol is also essential for resistance to oxidative, heat, and osmotic stress (Liu et al, 2017; Montañés et al., 2011; Hickman *et al.*, 2011; Kodedová and Sychrová, 2015). Therefore, metabolic engineering of mevalonate pathway for production of other isoprenoids will inevitably impact ergosterol homeostasis and stress resistance of the yeast cells. In *S. cerevisiae*, different kinds of stress stimuli trigger the same kind of transcriptional response where cluster of stress response genes are activated while housekeeping genes are down-regulated (de Nadal and Posas, 2015). Quick changes in genome-wide transcription is regulated through structural changes in the chromatin that are induced by the cooperation of chromatin remodeling complexes, histone variants, and covalent modifications of histones (Bannister and Kouzarides 2011. Our analysis of the carotenoid screen of the yeast gene deletion collection revealed that two chromatin remodeling complexes, SWR1 and INO80, have opposite effects on the isoprenoid production. In accordance with the roles of these complexes in Htz1 loading/unloading onto the chromatin, Htz1 deletion increased carotenoid production, suggesting a role for Htz1 in regulating isoprenoid metabolism.

The canonical histone Htz1 regulates gene transcription by preventing the spreading of the silent chromatin (Meneghini *et al.*, 2003) and facilitating the passage of RNA polymerase II (Santisteban *et al.*, 2011). Our analysis of genome-wide transcriptional changes in deletion of Htz1 and other chromatin regulatory factors revealed that a cluster of stress-response genes and cell wall biogenesis genes were differentially regulated in *htz1* and *set1* strains. This data was in agreement with the recent work that showed Htz1 could sense stress and regulate gene transcription accordingly (Liu *et al.*, 2015). Expression of most of these genes were also affected when yeast was engineered for high level production of carotenoids (Verwaal *et al.*, 2010). Authors of this study reasoned that transcription of stress-response and cell wall biogenesis genes were perturbed due to the high level of carotenoids in this strain. Another obvious consequence of over-production of carotenoids will be a decrease in ergosterol production, which is essential for the stability of cell membrane and growth under stressful conditions such as hypoxia, hyperosmolarity, and hypothermia. Hence, low levels of ergosterol might activate the stress-response genes in a Set1 and Htz1 dependent manner. This hypothesis is also supported by a recent work (South *et al.,* 2013) that shows Set1 dependent methylation is involved in maintaining the ergosterol homeostasis. Direct measurement of ergosterol levels in these strains confirmed this observation (Figure 5). Single gene deletions had only modest effect on ergosterol levels. However, *set1, htz1* double deletion strain accumulated ergosterol beyond homeostatic levels under stationary phase.

**Figure 5.**
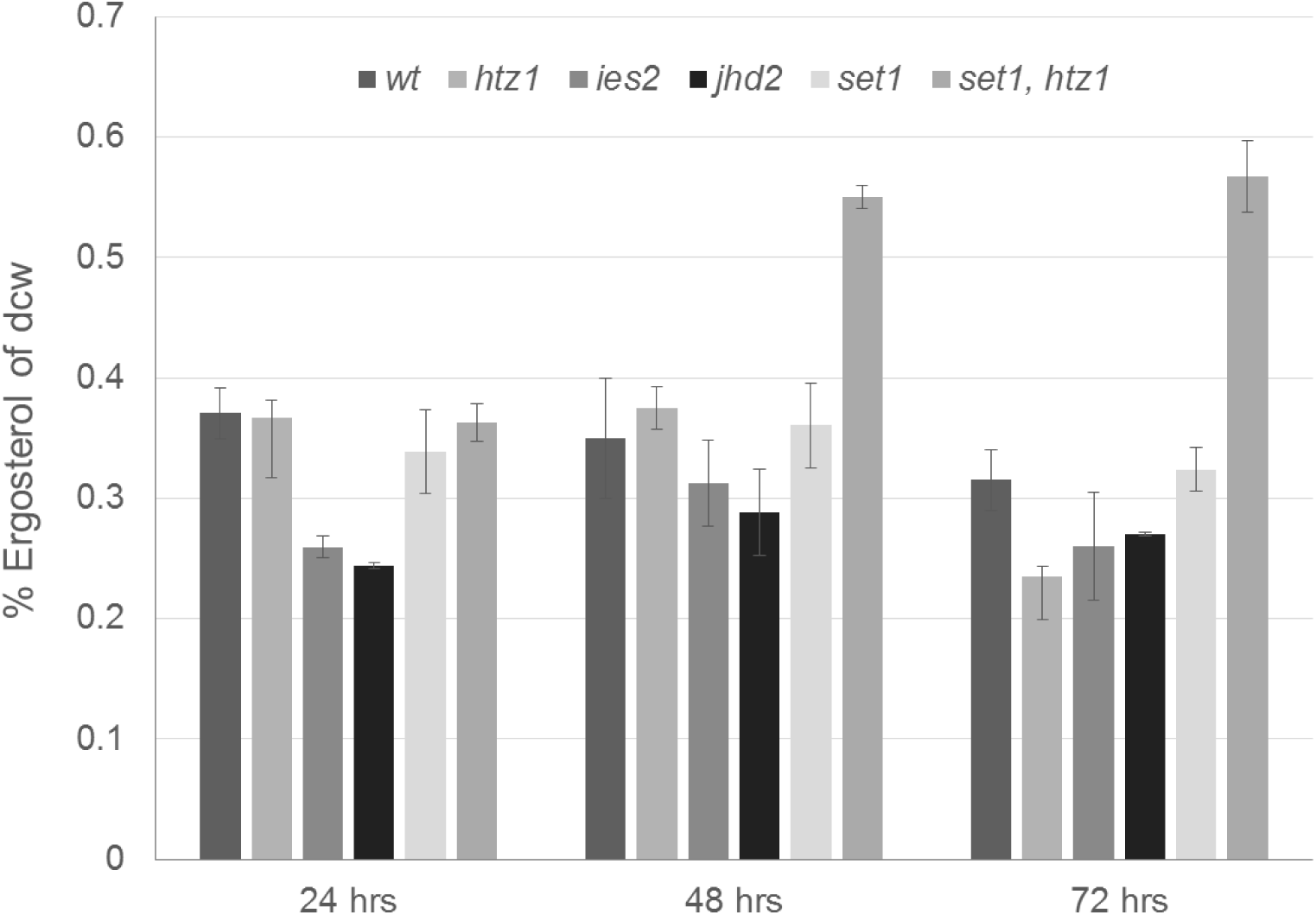
Total ergosterol content as percentage of dry cell weight (dcw) of wild-type and deletion strains (*htz1, ies2, jhd2, set1, htz1, set1*) in cultures harvested after 24, 48, and 72 hours of growth.

**Figure 6.**
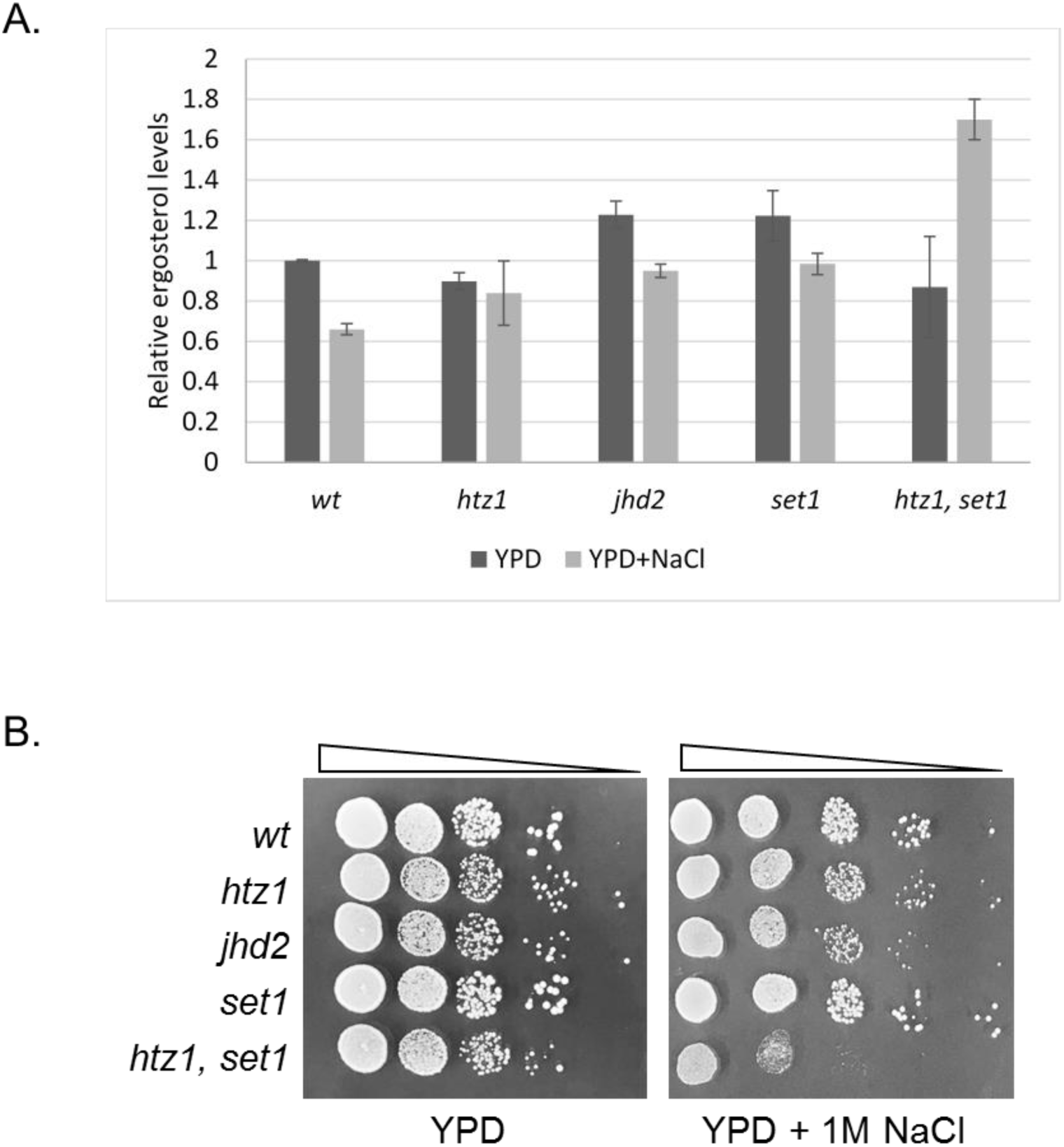
A. Relative ergosterol contents of strains grown under regular (YPD) and hyperosmotic (YPD+ 1M NaCl) conditions for 24 hrs. **B.** Overnight cultures grown in YPD were spotted in 10-fold serial dilutions onto regular (YPD) and hyperosmotic (YPD+1M NaCl) plates.

Under stationary phase stress, it is well known that trehalose and glycogen is upregulated (de Winde *et al.*, 1997). However, the *set1, htz1* strain failed to accumulate them, which is expected since ergosterol accumulation happens at the expense of these starches (Boulton and Quain, 2013). It is also well established that under hyperosmotic stress, ergosterol is down regulated and this down-regulation is essential for cell survival (Montañés *et al.*, 2011). The *set1, htz1* strain failed to down-regulate ergosterol under hyperosmotic conditions and resulted in growth defect, further confirming a role for these epigenetic factors in maintaining ergosterol homeostasis. This data was in accordance with recent findings of Nadal-Ribelles *et al.* (2015) on regulation of stress response genes through Set1 and SWR1C/Htz1 dependent chromatin remodeling.

Our work shows that eliminating Set1 and Htz1 dependent activation of stress response pathways results in failure to regulate ergosterol levels. Surprisingly, the relationship between ergosterol levels and stress response activation seems to be a two-way street where not only stress response activation affected the ergosterol levels but also the ergosterol levels activated stress response in a Set1/Htz1 dependent manner. One way for how ergosterol levels can induce Set1 dependent methylation could be via intracellular levels of SAM. Deletion of ergosterol synthesis genes result in accumulation of SAM (Shobayashi et al., 2006) and SAM levels dictate the extent of Set1 dependent histone methylation (Sadhu *et al.*, 2013). Follow-up experiments and further understanding of this cross-talk mechanism will not only help metabolic engineering efforts in isoprenoid production but also enable the medical community to find new means to regulate cholesterol levels in humans or treat fungal infections.

## Acknowledgements

This work was partially supported by the co-fund 2236 program of European Union and The Scientific and Technological Research Council of Turkey (TUBITAK) (Project number 114C014). We thank Oytun Eskiyenenturk for valuable advice and help in data analysis.

